# Cytokine-Mediated Degradation of the Transcription Factor ERG Impacts the Pulmonary Vascular Response to Systemic Inflammatory Challenge

**DOI:** 10.1101/2023.02.08.527788

**Authors:** Christopher M. Schafer, Silvia Martin-Almedina, Katarzyna Kurylowicz, Neil Dufton, Lourdes Osuna-Almagro, Meng-Ling Wu, Charmain F. Johnson, Aarti V. Shah, Dorian O. Haskard, Andrianna Buxton, Erika Willis, Kate Wheeler, Sean Turner, Magdalena Chlebicz, Rizaldy P. Scott, Susan Kovats, Audrey Cleuren, Graeme M. Birdsey, Anna M. Randi, Courtney T. Griffin

## Abstract

**Background:** During infectious diseases, pro-inflammatory cytokines transiently destabilize interactions between adjacent vascular endothelial cells (ECs) to facilitate the passage of immune molecules and cells into tissues. However, in the lung the resulting vascular hyperpermeability can lead to organ dysfunction. Previous work identified the transcription factor ERG as a master regulator of endothelial homeostasis. Here we investigate whether the sensitivity of pulmonary blood vessels to cytokine-induced destabilization is due to organotypic mechanisms affecting the ability of endothelial ERG to protect lung ECs from inflammatory injury.

**Methods:** Cytokine-dependent ubiquitination and proteasomal degradation of ERG was analyzed in cultured Human Umbilical Vein ECs (HUVECs). Systemic administration of TNFα or the bacterial cell wall component lipopolysaccharide (LPS) was used to cause a widespread inflammatory challenge in mice; ERG protein levels were assessed by immunoprecipitation, immunoblot, and immunofluorescence. Murine *Erg* deletion was genetically induced in ECs (*Erg*^*fl/fl*^*;Cdh5(PAC)Cre*^*ERT2*^), and multiple organs were analyzed by histology, immunostaining, and electron microscopy.

**Results:** In vitro, TNFα promoted the ubiquitination and degradation of ERG in HUVECs, which was blocked by the proteasomal inhibitor MG132. In vivo, systemic administration of TNFα or LPS resulted in a rapid and substantial degradation of ERG within lung ECs, but not ECs of the retina, heart, liver, or kidney. Pulmonary ERG was also downregulated in a murine model of influenza infection. *Erg*^*fl/fl*^*;Cdh5(PAC)-Cre*^*ERT2*^ mice spontaneously recapitulated aspects of inflammatory challenges, including lung-predominant vascular hyperpermeability, immune cell recruitment, and fibrosis. These phenotypes were associated with a lung-specific decrease in the expression of *Tek*, a gene target of ERG previously implicated in maintaining pulmonary vascular stability during inflammation.

**Conclusions:** Collectively, our data highlight a unique role for ERG in pulmonary vascular function. We propose that cytokine-induced ERG degradation and subsequent transcriptional changes in lung ECs play critical roles in the destabilization of pulmonary blood vessels during infectious diseases.

## INTRODUCTION

Vascular endothelial cells (ECs) form a dynamic, semi-permeable interface between blood and tissues. Therefore, ECs are among the first responders to systemic inflammatory challenges such as sepsis, a life-threatening condition in which bacterial infection of the blood triggers the release of abundant cytokines into the circulation^1,2^. In response to these stimuli, ECs adopt inflammatory phenotypes and participate in the initiation of local tissue responses^3,4^. By adjusting expression of cell surface proteins, activated ECs weaken their own intercellular junctions while also promoting interactions with circulating leukocytes^5^. Collectively, these alterations destabilize blood vessels, leading to vascular hyperpermeability that allows immune cell extravasation and the exchange of immune molecules between the blood and peripheral tissue. However, failure to resolve vascular inflammation in a timely manner results in excessive immune cell infiltration, pathological tissue edema, and ultimately organ dysfunction^6^.

For unclear reasons, blood vessels within the lung are particularly prone to pathological hyperpermeability in many inflammatory diseases^7–10^. During bacterial and viral infections, compromised vascular stability undermines pulmonary function by impairing efficient gas exchange between the dense capillary network and underlying airway epithelial cells^6,11–13^. This can result in acute respiratory distress syndrome (ARDS), which is associated with high mortality rates^8,11^. Since treatments that target the etiology of ARDS are currently lacking, patients are typically put on ventilators to support their reduced respiratory capacity^14^. Mounting experimental evidence suggests that stabilization of microvascular ECs may be a useful approach to alleviate pulmonary dysfunction^6,15–17^. Yet, refinement of these therapeutic strategies requires a deeper understanding of mechanisms responsible for the lung’s unique predisposition to vascular dysfunction under inflammatory challenges.

Here we demonstrate how organotypic regulation of the endothelial ETS family transcription factor ERG coordinates a vascular inflammatory response within the lung. ERG functions as a gatekeeper that suppresses vascular inflammation^18–23^. In turn, ERG is regulated by proinflammatory stimuli at the transcriptional and post-translational levels. In addition, recent reports demonstrate the capacity for proteolytic ERG degradation in cultured ECs, although the in vivo contexts for such degradation are not fully understood^24,25^. In this study, we demonstrate ERG ubiquitination and proteasomal degradation in ECs following stimulation with the proinflammatory cytokine TNFα in vitro and in vivo. Unexpectedly, we found that cytokine-induced ERG degradation in vivo occurs preferentially in pulmonary ECs. Downregulation of ERG is similarly seen in the lungs of mice challenged with a bacterial wall component or with viral infection. Importantly, genetic deletion of *Erg* in ECs throughout the body results in more severe vascular hyperpermeability, immune cell recruitment, and fibrosis in the lung than in other organs, suggesting that cytokine-induced ERG degradation may account for the lung’s unique susceptibility to inflammatory vascular dysfunction.

## METHODS

### Mice

Mice used in this study include C57Bl/6J (The Jackson Laboratory: #000664), *Il1r1*^*flox*^ (The Jackson Laboratory: #028398)^26^, *Tnfa*^*-/-*^ (The Jackson Laboratory: #003008)^27^, *Cdh5(PAC)CreERT2* (gift of Ralf Adams, Max Planck Institute for Molecular Biomedicine; available through Taconic: #13073)^28^,*VE-Cadherin-Cre* (The Jackson Laboratory: #006137)^29^, and *Tie2-Cre* (The Jackson Laboratory: #008863)^30^. EC-specific deletion of *Erg* was accomplished by crossing mice expressing *Cdh5(PAC)-CreERT2* with two independent *Erg*^*flox*^ mouse lines. For data presented in Fig. 5 and Fig. 6, *Erg*^*flox*^ mice were a gift of Joshua Wythe (Baylor College of Medicine; available through The Jackson Laboratory: #030988)^31^, and gene deletion was accomplished by oral gavage of 100 μL of 20 mg/mL tamoxifen (Sigma: #T5648) dissolved in peanut oil over three days. Mice were then returned to their cages for 3 – 4 weeks. For data presented in Fig. 7, *Erg*^*flox*^ mice were generated in our laboratory and described in detail elsewhere^32^; deletion of *Erg* was induced in adult mice, between 6–8 weeks, by tamoxifen injection (five injections of 0.5 mg daily). Genotyping for each allele was performed using the primers listed in Supplemental Table 2. All mice were maintained on a C57Bl/6J background. Mice were euthanized using isoflurane overdose (5%) followed by cardiac exsanguination and dissection. All studies using animals were performed in accordance with the NIH Guide for the Care and Use of Laboratory Animals and were approved by the Institutional Animal Care and Use Committees at the Oklahoma Medical Research Foundation or were conducted with ethical approval from Imperial College London under UK Home Office in compliance with the UK Animals (Scientific Procedures) Act of 1986.

### LPS, MG132, and pro-inflammatory cytokine treatment of mice

LPS, pro-inflammatory cytokines, and MG132 were administered to mice via intraperitoneal (LPS and MG132) or intravenous (LPS, TNFα, IL-1α, IL-1β, and IL-18) injection. Intraperitoneal LPS injection was performed using a 1-2 mg/mL stock solution of LPS (Sigma: #L3012; resuspended in sterile 0.9% saline) that was administered to mice at a dosage of 4 or 10 mg/kg BW for up to 72 h. Pre-treatment with MG132 was accomplished using a 20 mg/mL stock solution of MG132 (Enzo Life Sciences: #BML-PI102; resuspended in DMSO) that was diluted in 5% ethanol to a 0.5 mg/mL working stock, filter sterilized, and administered to mice at a dosage of 10 mg/kg BW for 3 h prior to LPS administration. Intravenous injections of LPS, TNFα (R&D Systems: #410-MT), IL-1α (Abcam: #ab256050), IL-1β (R&D Systems: #401-ML), or IL-18 (R&D Systems: #9139-IL) were performed via the retro-orbital sinus, as previously described^33^. Each compound was resuspended in sterile 0.9% saline from which 100-150 μL was injected into isoflurane-anesthetized mice at a final dosage of 1 mg/kg BW (LPS) or 50 μg/kg BW (TNFα, IL1α, IL-1β, and IL-18). Mice were then returned to their cages for 3 h prior to tissue collection.

### Influenza infection

Intratracheal influenza infection was performed on C57Bl/6J mice using a mouse-adapted A/Puerto Rico/8/1934 (PR8, H1N1) virus as previously described^34^. Mice were anesthetized by a single intraperitoneal injection of a Ketamine (60 mg/kg BW)/Xylazine (4.5 mg/kg BW) solution and then placed on an inclined surface to enable intratracheal administration of 30 μL of PBS containing either low (800 EID) or high (1050 EID) viral doses. Mice were then returned to their cages for 2 or 6 d, during which time they were monitored daily for signs of morbidity, such as weight loss.

### Immunoblot

For immunoblot analysis of murine tissues, mice were perfused with 10 mL warm PBS using a Bio-Rad Econo Pump (at a flow rate of 1 mL/min) and a 23 g needle inserted into the left ventricle of the heart. Lung, heart, liver, kidney, and retina tissues were then collected, washed in PBS, and snap frozen in liquid N_2_. Tissue samples were then lysed by sonication in RIPA buffer [50 mM Tris-HCl (pH 7.4), 150 mM NaCl, 1% SDS, 1mM EDTA, 0.5% sodium deoxycholate] with 1 mM phenylmethylsulfonyl fluoride and Protease Inhibitor Cocktail (Sigma: #P8340) added just before use. Tissue lysates were centrifuged at 10,000 X g for 10 min at 4°C to remove cell debris, and the protein concentrations of supernatants were determined using a BCA Protein Assay Kit (Thermo Scientific: #23227), following the manufacturer’s instructions. Samples were then diluted to 5-10 mg/mL in PAGE running buffer [62.5 mM Tris-HCl (pH 6.8), 2% SDS, 10% glycerol, 5% β-mercaptoethanol, 0.002% bromophenol blue] and heated at 95°C for 10 min. Protein samples (10-20 μg) were separated on 9% SDS-PAGE gels and then transferred to a PVDF membrane that was blocked for 1 h in 5% non-fat dry milk dissolved in TBST. Primary antibodies were diluted in 5% milk-TBST and incubated on membranes overnight at 4°C with gentle agitation. Membranes were then washed three times (15 min each) in TBST, and HRPconjugated secondary antibodies (diluted in 5% milk-TBST) were applied for 3 h at 25°C with gentle agitation. Secondary antibodies were detected using ECL Western Blotting Detection Reagent (GE Healthcare: #95038-562). Immunoblot band densitometry and normalization to GAPDH expression was performed using NIH ImageJ software^35^.

### Tissue section imaging by immunofluorescence and immunohistochemistry

To generate tissue sections for immunofluorescence and immunohistochemistry, mice were euthanized and perfused via the heart’s left ventricle with 10 mL warm PBS followed by 5 mL of 4% paraformaldehyde to fix tissues. To inflate the lung, 2-3 mL of a 10% solution of buffered formalin was administered using a 26 g needle inserted into the trachea. Once inflated, the trachea was sealed using a surgical suture. Lungs were then excised and incubated overnight in 10% formalin. Paraffin-embedded sections were primarily used for immunohistochemistry by hematoxylin/eosin, Masson’s trichrome and DAB staining. OCT-embedded sections were prepared by sequential 1 h incubations in 10%, 15%, and 20% sucrose-PBS and then incubated overnight at 4°C in a 1:1 solution of 20% sucrose and OCT compound (Fisher Scientific: #23730571). The next day, tissues were embedded with OCT compound in cryomolds, and tissue sections (5-10 μm) were collected on an HM525 NX cryotome (Epredia). Tissue sections were dried for 30 min at 25°C, washed three times in PBS, permeabilized for 20 min in 0.2% Triton X-100, and blocked overnight at 4°C in 10% donkey serum (Jackson ImmunoResearch Lab: #102644-006), 3% BSA-PBS. Antibodies were diluted in 0.02% Triton X-100, 1% BSA-PBS and applied to sections overnight at 4°C and 3 h at 25°C, respectively. For studies in influenza-infected mice, immunofluorescence was performed using an antibody directed against hemagglutinin that was obtained through the NIH Biodefense and Emerging Infections Research Resources Repository, NIAID, NIH: Polyclonal Anti-Influenza Virus H1 (H0(Hemagglutinin (HA), A/Puerto Rico/8/34 (H1N1), (antiserum, goat), NR-3148). Images were collecting using a Nikon TiE Eclipse epifluorescence microscope or a Nikon C2 confocal microscope and analyzed using Nikon Elements software (v4).

### Tissue permeability assays

For Evans blue permeability assays, mice were anesthetized by isoflurane inhalation, and 100150 μL of 1% Evans blue (Sigma: #E2129) that was filtered through a 0.2 μm membrane was administered to mice via the retro-orbital sinus at a final dosage of 40 mg/kg BW. After 1 h incubation, mice were euthanized and perfused with 10 mL warm PBS. Lung, heart, liver, and kidney tissues were then excised, washed in PBS, and dried overnight at 37°C. The next day, dry tissue weights were recorded, and 500 μL formamide was added to each sample followed by a 48 h incubation at 55°C to extract the Evans blue dye. Samples were briefly centrifuged to remove tissue fragments, and supernatants were used to quantity Evans blue spectrophotometrically (OD_670_) by comparison to a standard curve. Final values were normalized to tissue dry weight. Intravenous injection of FITC-Dextran (MW=150,000; Millipore Sigma: FD150S) was similarly performed via the retro-orbital sinus. Mice were administered 100-150 μL of a filtered 25 mg/mL FITC-Dextran solution at a final dosage of 100 mg/kg BW. FITC-Dextran was allowed to circulate for 1 h, at which time mice were euthanized, perfused with warm PBS, fixed in 4% paraformaldehyde, and then embedded in OCT for tissue sectioning.

### Real-time quantitative reverse transcription PCR (qPCR)

Total RNA was extracted from murine tissue samples or cultured ECs using Trizol and following the manufacturer’s instructions. RNA (1 μg) was then converted to cDNA using an iScript cDNA Synthesis Kit (Bio-Rad: #1708891BUN), which was then analyzed on a BioRad CFX96 Real Time Thermocycler using 2X SYBR green qPCR master mix (Life Technologies: #4312704). Target gene expression was normalized against three reference genes: *Actb, Gapdh*, and *Rn18s* (**Supplemental Table 2**).

### EC culture and treatments

HUVECs, isolated as previously described^36^ or purchased from commercially available sources (ATCC: #PCS-100-010), were cultured in complete EGM-2 media (Lonza: CC-33162). For TNFα treatment of HUVECs, cells were treated with 10 ng/mL TNFα (R&D Systems: #410-MT) for up to 10 h followed by the isolation of RNA and protein for qPCR and immunoblot, respectively. For MG132 treatments, HUVECs were treated with 10 μM MG132 (Enzo Life Sciences: #BML-PI102) or vehicle (DMSO) for 30 min prior to a 6 h TNFα treatment. Immunoprecipitation of endogenous ERG was performed using HUVECs following MG132/TNFα treatment. Cells were harvested in lysis buffer supplemented with protease and phosphatase inhibitors and the isopeptidase inhibitor N-ethylmaleimide (NEM) (Sigma: 04259). After clarification by centrifugation, 1 mg of total protein cell lysate was incubated with 4 µg of rabbit anti-human-ERG Ab (Santa Cruz: #sc-28680) overnight at 4°C. The immuno-complexes were precipitated with True-blot goat-anti-rabbit Ig beads (Rockland: #00-8800-25) for 4 hours at 4°C and detected by western blot with anti-ERG and anti-ubiquitin antibodies.

For siRNA-mediated *Erg* gene knockdown, HUVECs were grown to ∼80% confluence followed by transfection with 100 nM human ERG Silencer Select (ID #s4811) or nontargeting siRNA oligos (Life Technologies: ERG #4392420 and nontargeting #AM4635) using lipofectamine RNAiMAX (Life Technologies: #13778150) in serum-free OptiMEM (Life Technologies #31985070). Gene knockdown proceeded for 48 h prior to cell collection in Trizol for subsequent RNA isolation and qPCR analysis.

### Isolation and cloning of Erg wildtype and mutant Myc-tagged cDNA

HUVECs were grown on 1% gelatin-coated plates in EGM-2 medium (Lonza: CC3156) for 48 hrs. Total RNA was extracted using the RNeasy kit (Qiagen: #74106). First strand cDNA synthesis was performed using 1 µg total RNA with Superscript III reverse transcriptase (Invitrogen: #18080093). PCR amplification was performed with 1 µl cDNA and 0.5 µM Erg001-F forward and Erg-R-Myc reverse primers (**Supplemental Table 2**). PCR products were purified using the QIAquick gel extraction kit (Qiagen: #28706X4) and ligated into pcDNA3.1 (Invitrogen: V79020). All clones were sequenced from both ends. The pcDNA3.1 ERG (GenBank accession. NM_182918) Myc-tagged construct was mutated using the QuickChange Lightning Multi SiteDirected Mutagenesis kit (Agilent: #210513). All primers were designed using the QuickChange Primer Design Program (Agilent) and are listed in Supplemental Table 2. Briefly, pcDNA3.1 ERG-Myc-tagged plasmid was amplified using one or several primers designed to mutate a specific triplet that codifies from lysine to arginine. All constructs were verified by DNA sequencing. The different mutants generated were: K67R, K89R, K92R, K111R, K271R, K282R, K67-111R, K271-282R and K67-282R.

For analysis of ubiquitination of transfected ERG Myc-tagged, HeLa cells were transfected with ERG Myc WT or K/R mutants and Flag-tagged ubiquitin (kindly provided by Prof. H. Walczak) or empty vectors. Transfection was carried out with GeneJuice (Novagen: #C134443). After 24 hours, cells were pretreated with vehicle (DMSO) or with MG-132 for 6 hours. Incubation of cell lysates with polyubiquitin-affinity beads (Calbiochem: #662200-1KIT) was carried out according to the manufacturer’s protocol followed by immunoblotting with an anti-ERG Ab (Santa Cruz: #sc-353).

### Analysis of ERG protein stability in HeLa cells

HeLa cells were transfected with ERG Myc WT or ERG Myc K67-282R mutant. After 8 hours, the transfected cells were treated with cycloheximide (Sigma: C7698) at a final concentration of 100 µg/ml for 0, 4, and 6 hours in the presence of MG-132 (10 µM; Calbiochem) or DMSO (vehicle). Cells were harvested in lysis buffer supplemented with protease inhibitors, and lysates were subjected to immunoblot analysis to assess the expression level of ERG protein (Santa Cruz: #sc-353). GAPDH was used as internal control (Millipore: #MAB374).

### Transmission electron microscopy

Lung and kidney tissue from *Erg*^*iECko*^ and control mice were immersion fixed at room temperature in 0.1 M cacodylate buffer (pH 7.5) containing 4% formaldehyde and 2% glutaraldehyde followed by a second fixation in 1% OsO_4_. After dehydration, fixed specimens were embedded in expoy resin. Ultrathin resin sections were stained with solutions containing uranyl acetate and lead acetate and were visualized on an FEI Tecnai G2 transmission electron microscope.

### Statistics

Prism 7.0 software (GraphPad) was used for all statistical assessments. Statistical significance between two groups was assessed by unpaired parametric two-tailed t-tests. Comparison of multiple means was made using repeated-measures ANOVA with multiple comparisons between individual group means. Statistically analyzed data are presented as mean ± SD.

### Data Availability

The data underlying this article are available in the article and in its online supplementary material.

## RESULTS

### TNFα induces ERG ubiquitination and degradation in cultured ECs

We and others have reported that stimulation of cultured ECs with the pro-inflammatory cytokine TNFα leads to ERG downregulation in vitro^20,37^. To understand the mechanism of TNFα-induced ERG downregulation we treated Human Umbilical Vein ECs (HUVECs) with the cytokine and assessed ERG protein and transcript expression by immunoblot and qPCR, respectively. TNFα stimulation led to ERG protein downregulation in HUVECs **(Fig. 1A)** that was not accompanied by a similar decrease in *Erg* transcripts **(Supplemental Fig. 1A)**, suggesting a posttranscriptional mechanism of ERG downregulation, such as regulated protein degradation. In support of this hypothesis, pre-treatment of HUVECs with MG312, an inhibitor of the proteasome, prevented TNFα-induced ERG downregulation **(Fig. 1B** and **Supplemental Fig. 1B)**. Since proteins are commonly poly-ubiquitinated prior to degradation^38^, we next assessed the ubiquitination status of ERG in HUVECs after TNFα treatment. HUVECs were treated with TNFα in the presence or absence of MG132, and cell lysates were subjected to ERG immunoprecipitation and probed for ubiquitin by immunoblot. TNFα treatment led to the appearance of high molecular weight, poly-ubiquitinated ERG isoforms that increased in abundance following proteasomal inhibition by MG132 **(Fig. 1C)**.

**FIGURE 1.**
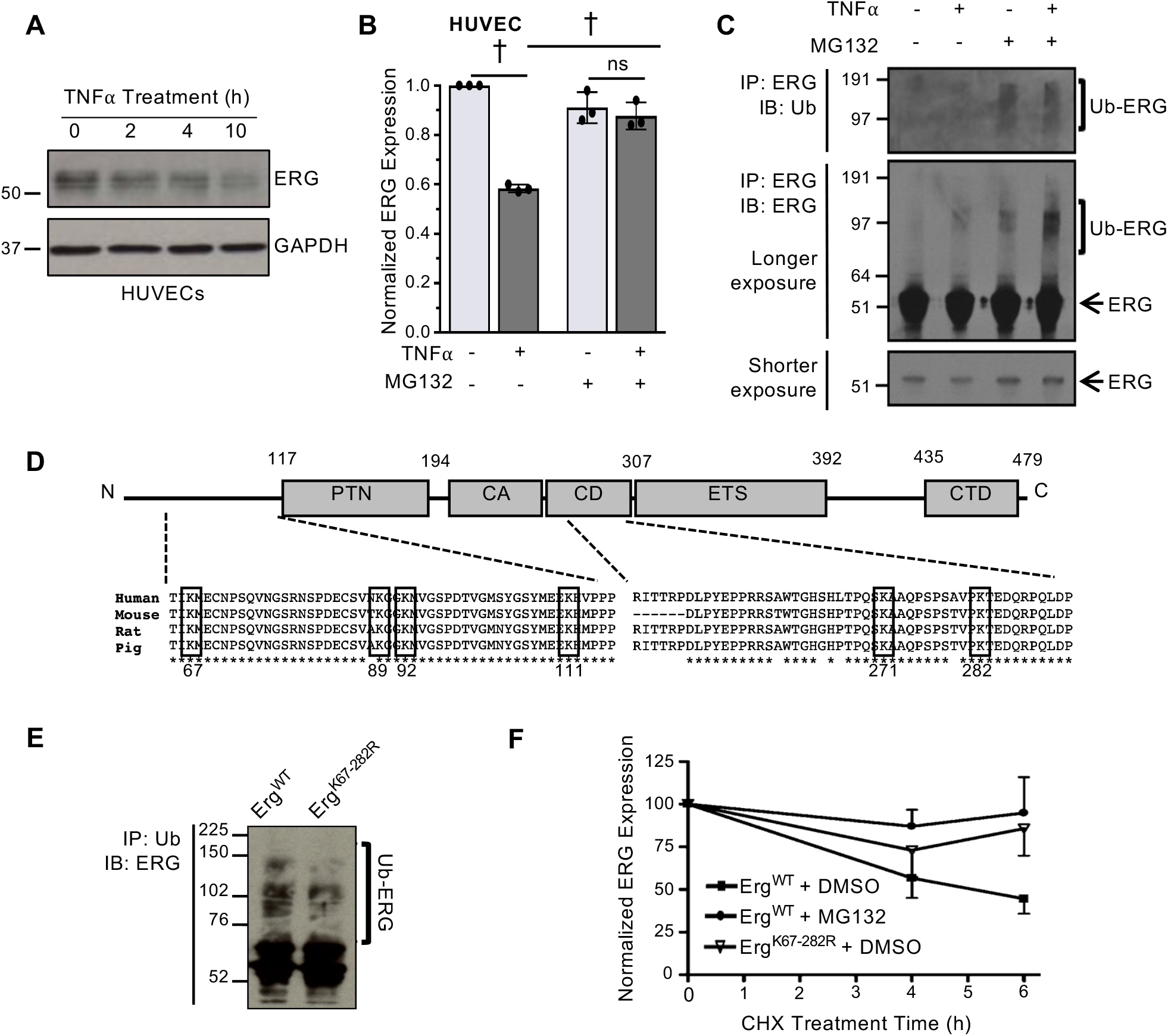
The p*ro-inflammatory cytokine TNF*α *promotes ERG ubiquitination and proteolytic degradation in vitro*. **(A)** TNFα (10 ng/mL) was applied directly to cultured HUVECs, and ERG expression was assessed by immunoblot at the indicated times. **(B)** TNFα (10 ng/mL for 6 h)induced ERG downregulation in HUVECs was assessed by immunoblot following a 30 min pretreatment with MG132 (10μM). **(C)** Following a similar TNFα/MG132 treatment, ERG was immunoprecipitated from HUVECs, and samples were immunoblotted for ERG and ubiquitin (Ub). Longer exposure of the ERG immunoblot revealed the presence of high molecular weight, ubiquitinated ERG isoforms. **(D)** Alignment of human, mouse, rat, and pig ERG sequences highlighting six conserved lysine residues (K) predicted as sites of ubiquitination. **(E, F)** Mutation of all putative sites to arginine (Erg^K67-282R^) reduced the presence of ubiquitinated ERG isoforms (G) and prolonged ERG half-life (H) when constructs were expressed in HeLa cells. ^†^p<0.05 (twoway ANOVA)

Using the ubiquitination site prediction software UbPred, we identified six putative ubiquitination sites within ERG: K67, K89, K92, K111, K271 and K282 **(Fig. 1D)**. To determine if any of these sites are ubiquitinated, we transiently transfected HeLa cells with Myc-tagged wildtype ERG or with ERG mutants in which the putative ubiquitinated lysine (K) residues had been mutated to arginine (R) **(Supplemental Fig. 1C)**. Treatment of transfected HeLa cells with MG132 for 6 h led to the accumulation of poly-ubiquitinated ERG isoforms, which could be further enhanced by co-transfection with a FLAG-tagged ubiquitin construct **(Supplemental Fig. 1D)**. We found that individual mutations of K67R, K271R and K282R each resulted in a partial reduction in ERG ubiquitination **(Supplemental Fig. 1E)**. However, mutation of all six lysine residues (K67-282R) had the most profound effect on ERG ubiquitination **(Fig. 1E** and **Supplemental Fig. 1E)** and prolonged the half-life of ERG protein following cycloheximide treatment **(Fig. 1F)**.

### Inflammatory stimulation promotes lung-specific ERG downregulation in vivo

Next, we asked if TNFα stimulation was similarly capable of driving ERG degradation in vivo. We performed intravenous injections of TNFα into wildtype mice and used immunoblot to assess ERG expression in two highly vascularized tissues, the lung and heart. Three hours after challenge we observed a substantial TNFα-induced downregulation of ERG in the lung but not the heart **(Fig. 2A-B)**. Moreover, a similar pattern of lung-specific ERG downregulation was observed following treatment with the pro-inflammatory cytokines IL-1α and IL-1β, but not IL-18 **(Fig. 2A-B** and **Supplemental Fig. 2A)**.

**FIGURE 2.**
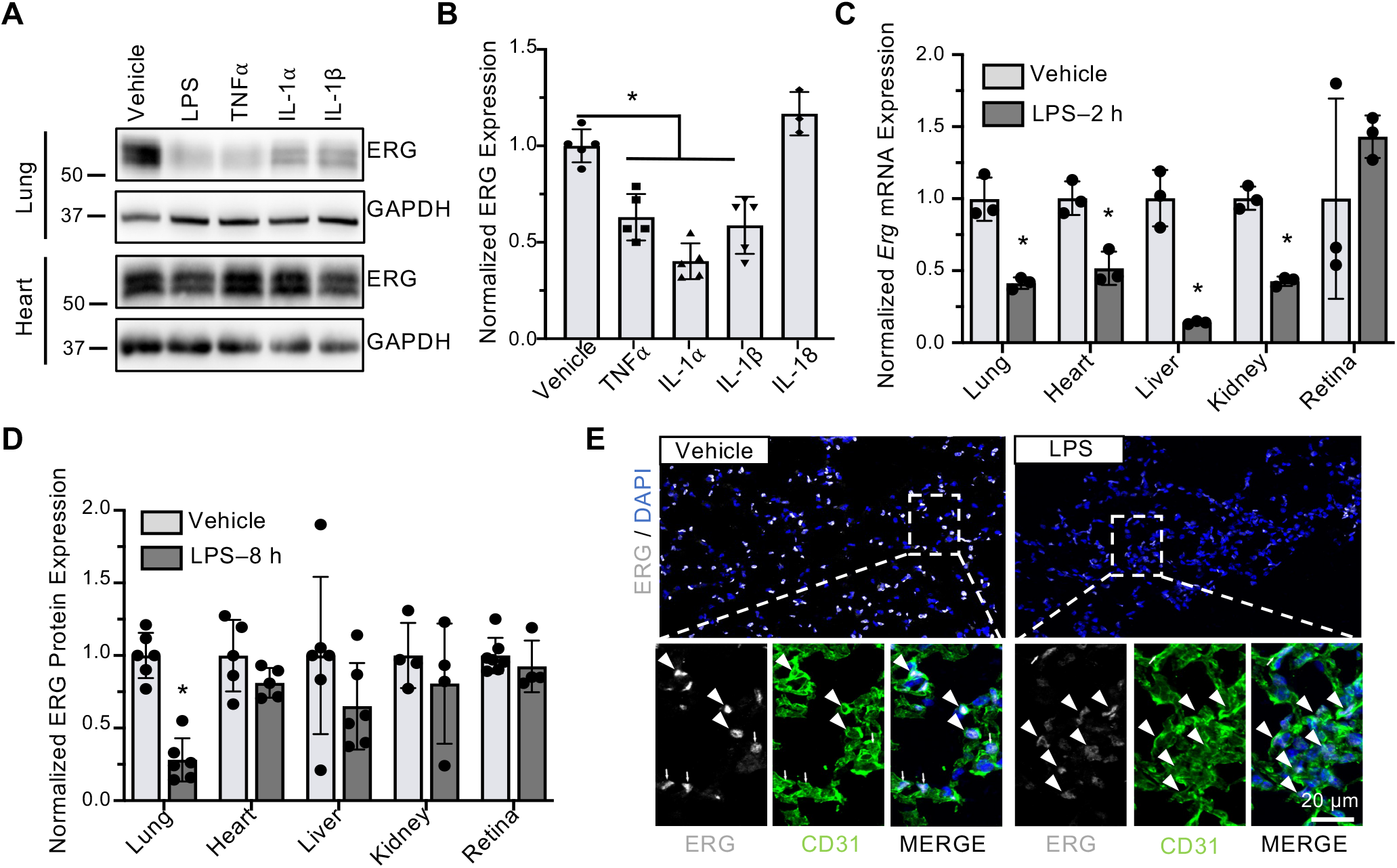
Pro-inflammatory stimuli promote lung-specific ERG protein downregulation. **(A, B)** ERG expression was assessed by immunoblot (A) and quantified (n=4-5) (B) in the lung and heart at 3 h after intravenous injection of LPS (1 mg/kg BW), TNFα, IL-1α, IL-1β, or IL-18 (50 µg/kg BW for each cytokine). **(C, D)** WT mice were administered intraperitoneal injections of LPS (4 mg/kg BW) or vehicle (0.9% NaCl). *Erg* transcript expression was quantified by qPCR in the indicated organs at 2 h after LPS exposure (n=3) (C), and ERG protein expression was quantified by immunoblot at 8 h after LPS exposure (n=4-6) (D). **(E)** Expression of ERG (grey) in CD31^+^ (green) ECs (white arrows) within the lung was assessed by immunostaining tissue sections collected 8 h after LPS exposure. DAPI (blue) was used as a nuclear counterstain. *p<0.05 (two-way t-test)

Pulmonary ECs are particularly prone to activation during systemic inflammatory challenges, such as bacterial sepsis, in which circulating pro-inflammatory cytokines like TNFα are upregulated. We therefore asked if lung-predominant ERG degradation occurs during a murine model of LPS-induced sepsis. Acute administration of LPS (4 mg/kg BW) to adult wildtype mice resulted in the transcriptional downregulation of *Erg* within the lung, heart, liver, and kidney after 2 h, while the immune-privileged retina was spared from *Erg* downregulation within this timeframe **(Fig. 2C)**. However, at 8 h post-injection, significant LPS-induced downregulation of ERG protein was only observed in the lung by immunoblotting **(Fig. 2D)**. This was confirmed by immunostaining for ERG in tissue sections at 8 h post-LPS challenge, revealing a loss of ERG expression within CD31^+^ capillary ECs of the lung **(Fig. 2E)** that was not observed in the heart, kidney, or liver **(Supplemental Fig. 3A-D)**. Furthermore, at this same LPS dose pulmonary ERG expression began to recover by 24 h, though we continued to observe a slight but significant reduction in pulmonary ERG expression up to 48 h post-injection **(Supplemental Fig. 2B-C)**. A similar pattern of protein downregulation in the lung following LPS challenge was also observed for the transcription factor FLI1 **(Supplemental Fig. 2D)**, a homologue of ERG with overlapping and compensatory functions^22,39^.

### LPS administration leads to cytokine-induced ERG degradation in the lung

We then used genetic models to determine which specific pro-inflammatory cytokine(s) cause LPS-induced ERG downregulation in the lung. When we administered LPS (4 mg/kg BW) to mice in which the TNFα gene was globally deleted (*Tnfa*^*-/-*^), we saw a significant rescue of the LPS-induced pulmonary ERG downregulation seen in littermate control mice **(Fig. 3A)**. Global deletion of the primary receptor for IL-1α/β (*Il1r1*^*flox/flox*^*;Sox2-Cre*^*+*^ = *Il1r1*^*Gko*^) was also sufficient to prevent LPS-induced downregulation of pulmonary ERG **(Fig. 3B)**. However, deletion of *Il1r1* specifically from ECs using two independent Cre lines failed to rescue LPS-induced ERG downregulation in the lung **(Fig. 3C** and **Supplemental Fig. 2E)**, suggesting that IL-1α/β likely works through an intermediate cell type to stimulate pulmonary endothelial ERG downregulation.

**FIGURE 3.**
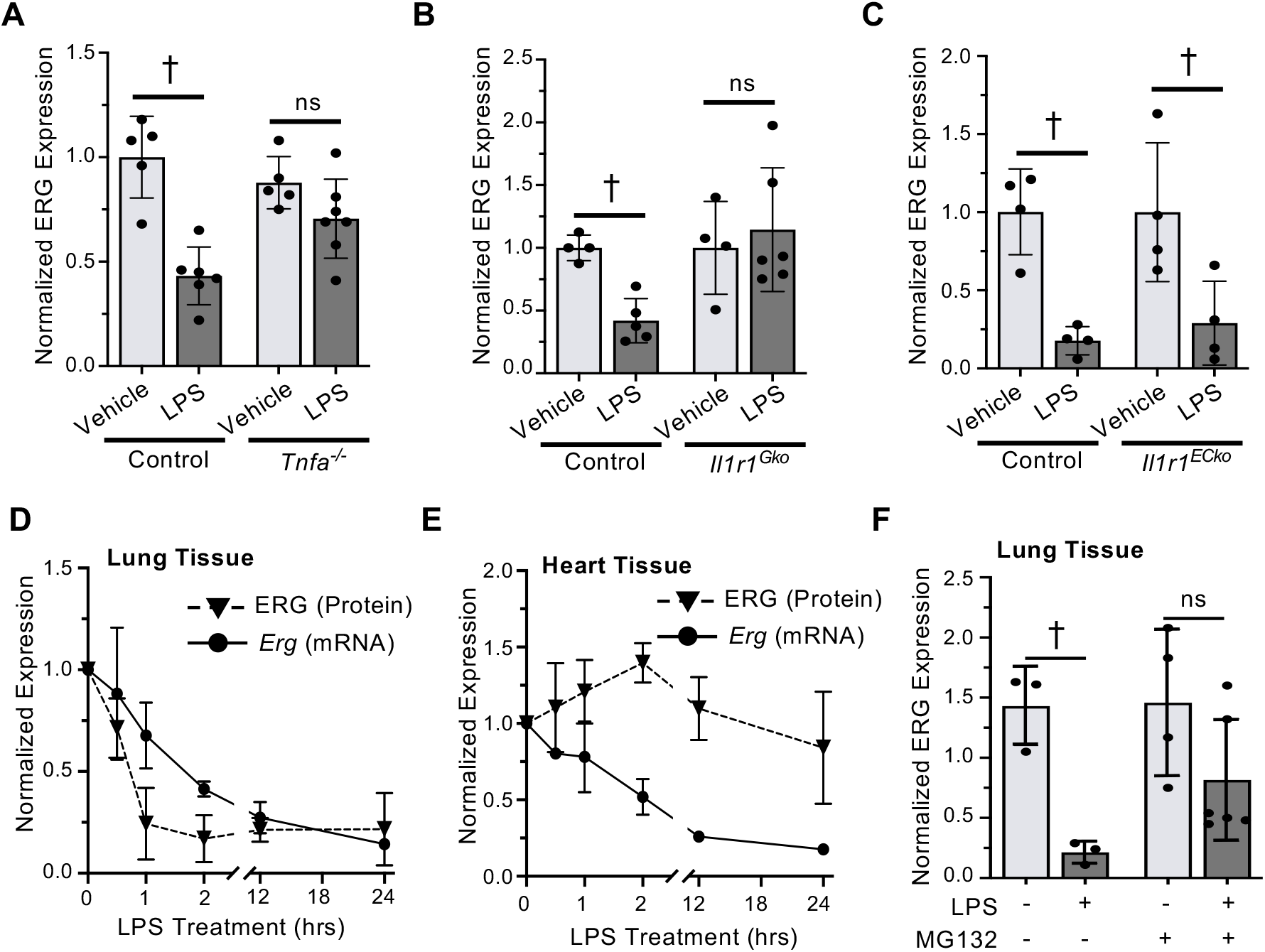
Pro-inflammatory cytokines promote LPS-induced ERG degradation in the lung. **(A - C)** ERG expression was assessed by immunoblotting lung lysates at 4 h after intraperitoneal administration of LPS (4 mg/kg BW) in mice following global deletion of *Tnfa* (*Tnfa*^*-/-*^) (n=5-6) (A), global deletion of *Il1r1* (*Il1r1*^*Gko*^) (n=4-5) (B), OR EC-specific deletion of *Il1r1* (*Il1r1*^*ECko*^) (n=5) (C). **(D, E)** Wildtype mice were given an intraperitoneal injection of LPS (10 mg/kg BW). At 0.5-24 h after LPS challenge, ERG protein and transcript expression were quantified in lung (D) and heart (E) tissues by immunoblot and qPCR, respectively (n=3-4). **(F)** LPS (4 mg/kg BW)- induced ERG downregulation in the lung was assessed by immunoblot following a 3 h pretreatment with vehicle or MG132 (10 mg/kg BW) (n=3-6). ^†^p<0.05 (two-way ANOVA)

We next monitored the kinetics of pulmonary ERG downregulation following acute LPS (10 mg/kg BW) challenge. LPS-induced ERG protein downregulation in the lung was rapid: within 1 h of LPS administration, we observed a >70% reduction in ERG protein expression, which is consistent with a half-life of ∼0.5 h **(Fig. 3D)**. In contrast, cardiac ERG protein expression was unaffected by LPS challenge during this time frame and only began to trend downwards by 24 h post-LPS challenge **(Fig. 3E)**. Interestingly, we observed LPS-induced transcriptional downregulation of *Erg* in both the lung and heart. However, transcriptional downregulation of *Erg* in the lung lagged behind the downregulation of ERG protein **(Fig. 3D)**, which is consistent with LPS-induced ERG degradation in vivo similar to what was observed for HUVECs in vitro **(Fig. 1B)**. In support of this hypothesis, pre-treatment of wildtype mice with MG132 partially prevented LPS-induced ERG downregulation in the lung **(Fig. 3F)**

These data suggest a lung-specific mechanism of cytokine-induced ERG ubiquitination and degradation. We therefore asked if lung ECs might uniquely express a ubiquitin ligase that targets ERG for degradation. We took advantage a recently published list of EC-expressed gene transcripts within the lung, heart, liver, and kidney that was assembled using translating ribosome affinity purification (EC-TRAP)^40^. This list was cross referenced against a comprehensive database of mammalian ubiquitin ligase genes^41^ to generate a list of ∼280 ubiquitin ligases expressed within ECs of each organ **(Supplemental Table 1)**. Included in this list were previously identified regulators of ERG ubiquitination such as *Trim25*, which impacts ERG degradation in ECs, and *Spop*, which can degrade ERG in prostate cancer cells^25,42,43^. Notably, *Trim25* was nearly two-fold more abundant in lung ECs relative to ECs of the kidney and liver, although lung and heart ECs had more comparable levels of *Trim25* expression **(Supplemental Table 1)**. Moreover, for an additional ∼30 ubiquitin ligase genes, we observed greater than two-fold higher expression in lung ECs compared to the ECs from any other organ **(Supplemental Table 1)**. However, future studies will be needed to determine if expression of these ubiquitin ligases play a role in lung-specific ERG degradation.

### ERG is downregulated in pulmonary ECs after an influenza challenge

Since pro-inflammatory cytokines, such as TNFα, are also produced in the lung during viral infections, we determined if pulmonary ECs downregulate ERG during a murine model of influenza infection^44^. We assessed ERG expression following intratracheal exposure to a mouse-adapted influenza virus and observed a significant decrease in ERG protein expression 6 d after administration of low [800 egg infectious dose (EID)] and high (1050 EID) viral doses **(Fig. 4A)**. Immunostaining of influenza-infected lungs revealed reduced ERG expression relative to the lungs of vehicle-injected mice **(Fig. 4B)**. Intriguingly, ERG downregulation was more substantial in pulmonary capillary beds relative to larger caliber blood vessels, which appeared to maintain higher ERG expression **(Fig. 4B** and **Supplemental Fig. 4)**. In addition, the influenza virus was regionally localized within infected lungs, as evidenced by immunostaining for the influenza protein hemagglutinin (HA) **(Fig. 4C)**. Importantly, immunostaining of influenza-infected lungs revealed a loss of ERG expression that co-localized with the heavily infected (HA-positive) regions of the lung **(Fig. 4C)**.

**FIGURE 4.**
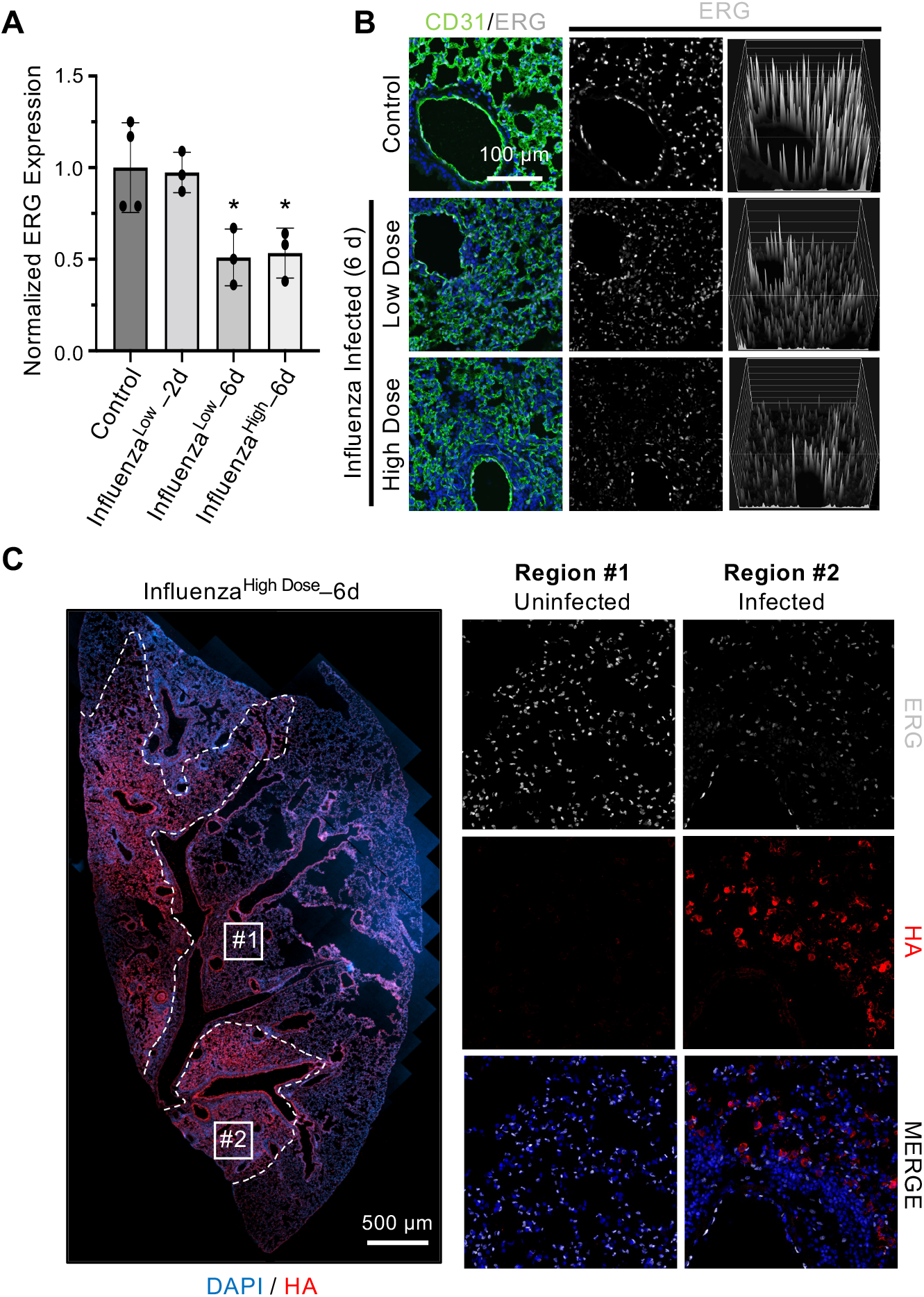
ERG is downregulated in the lung during pulmonary influenza infection. **(A)** ERG expression was quantified by immunoblot using murine lung tissue collected 2 and 6 d after low (800 EID) and high (1050 EID) doses of influenza were administered via the trachea (n=3). **(B)** Expression of ERG (grey) in ECs (CD31; green) was assessed by immunostaining lung tissue sections collected from control or influenza-infected mice. Reduced ERG expression within pulmonary capillary ECs (but not within ECs of larger caliber vessels) of influenza-infected mice is visualized by a surface intensity representation of the ERG channel (right panels); DAPI (blue) was used as a nuclear counterstain. **(C)** Immunostaining for the influenza hemagglutinin protein (HA; red) was used to identify uninfected (#1) and infected (#2) regions of a lung section. Reduced expression of ERG (grey) was observed in infected regions relative to uninfected regions of the same lung section. *p<0.05 (two-way t-test)

### Transcriptional regulation of *Tek*/TIE2 by ERG in pulmonary ECs in vivo

ERG is a master regulator of endothelial gene expression^19,45^; thus, the downregulation of ERG during a systemic inflammatory challenge likely has a substantial impact on pulmonary EC transcripts. One of the key mediators of lung vascular homeostasis regulated by ERG is TIE2 (gene name = *TEK*)^18,23^, an EC-expressed receptor tyrosine kinase that maintains vascular stability under homeostatic conditions^46^. The connection between *TEK*/TIE2 repression and pulmonary vascular dysfunction in sepsis/ARDS is well documented^47–51^. During inflammation, multiple pathways contribute to the repression of TIE2 expression and activity, resulting in pulmonary blood vessel destabilization^52^. However, little is known regarding the mechanism(s) of transcriptional *TEK* repression in the inflamed lung.

In agreement with our previous work^18^, siRNA-mediated *ERG* knockdown in HUVECs led to a downregulation of *TEK* transcript expression **(Fig. 5A)**. We therefore used HUVECs to investigate the molecular mechanisms through which ERG regulates *TEK* expression. Analysis of ERG binding to DNA in HUVECs via chromatin immunoprecipitation-sequencing (ChIP-seq)^45^ identified an ERG binding peak associated with an intronic region mapped to the *TEK* gene locus, downstream of the transcription start site (TSS) **(Fig. 5B)**. The site was also enriched for the histone modification H3K27Ac and for RNA polymerase II (RNAPol2), suggesting the presence of an active enhancer region. ChIP-qPCR confirmed ERG enrichment at the *TEK* intronic region **(Fig. 5C)**.

**FIGURE 5.**
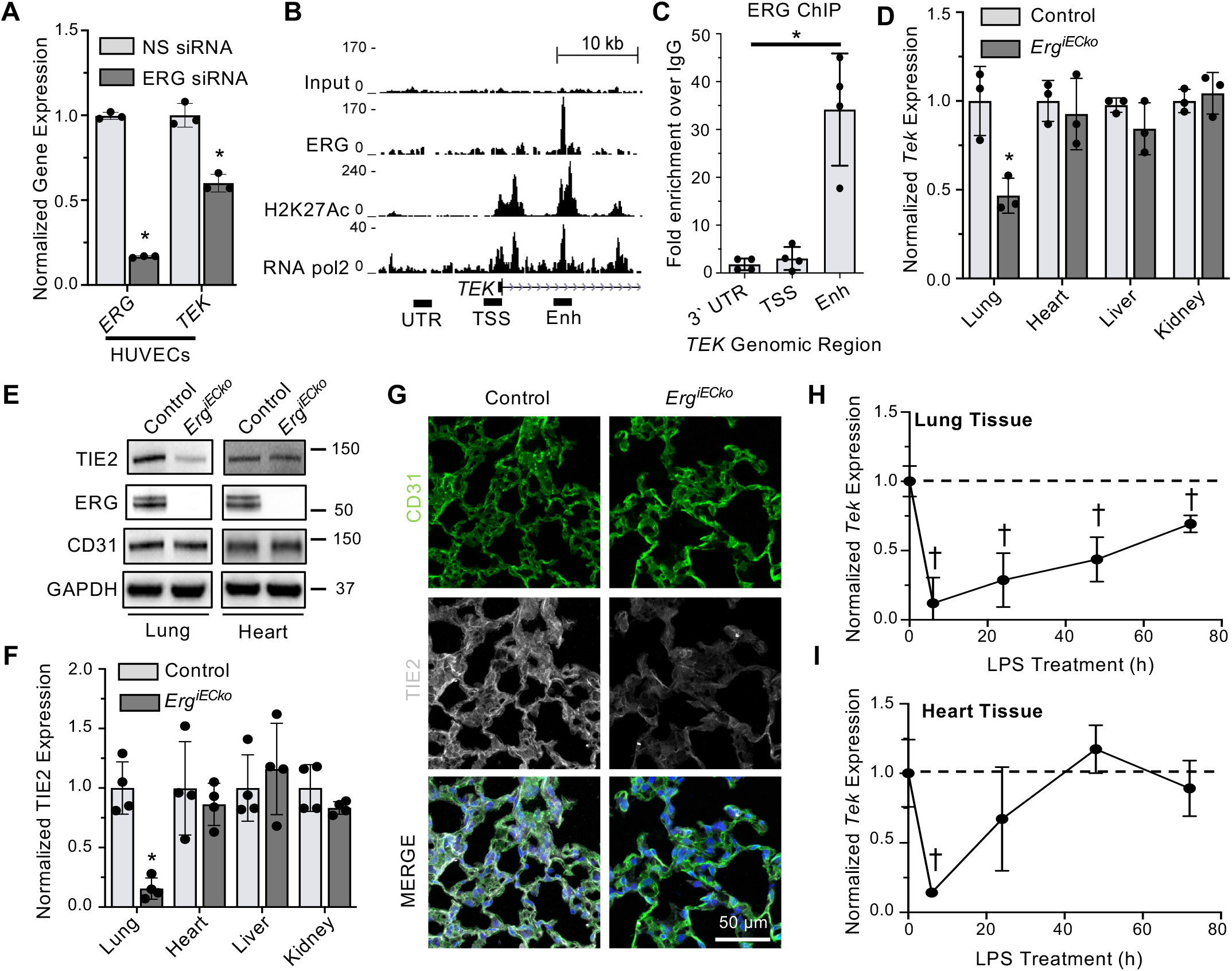
Tek is transcriptionally regulated by ERG. **(A)** *TEK* and *ERG* expression in HUVECs was assessed by qPCR following siRNA-mediated *ERG* knockdown. **(B**,**C)** ERG ChIP-seq (B) identified occupancy of ERG at the *TEK* transcriptional start site (TSS) as well as an intragenic region that co-localizes with H3K27Ac, a marker of active enhancers, in HUVECs. ERG binding to the TSS an enhancer region was confirmed by ChIP-qPCR (C). **(D)** *Tek* transcript expression was assessed by qPCR in the lung, heart, liver, and kidney of *Erg*^*iECko*^ and control littermates (n=3). **(E, F)** TIE2 protein expression in *Erg*^*iECko*^ and control littermates was assessed by immunoblot (E) and quantified (n=4) in the lung, heart, liver, and kidney (F). **(G)** Expression of TIE2 (grey) in CD31^+^ (green) ECs in the lungs of control and *Erg*^iECko^ mice was assessed by immunofluorescence. DAPI (blue) was used as a nuclear counterstain. **(H**,**I)** *Tek* expression was assessed by qPCR in WT murine lung (H) and heart (I) tissues collected 6-72 h after LPS (4 mg/kg BW) challenge (n=5-7). ^†^p<0.05 (one-way ANOVA) *p<0.05 (two-way t-test)

To investigate whether ERG regulates the expression of *Tek*/TIE2 *in vivo*, we genetically deleted *Erg* from ECs using a tamoxifen-inducible EC-specific Cre^28^ **(Supplemental Fig. 5A)** and a floxed allele of *Erg* [*Erg*^*fl/fl*^*;Cdh5(PAC)-CreERT2* = *Erg*^*iECko*^], and we used qPCR to quantify *Tek* transcripts in the lung, heart, liver, and kidney. Surprisingly, we observed significant *Tek* downregulation in the lungs of *Erg*^*iECko*^ mice but not in the other organs tested **(Fig. 5D)**. A similar pattern was observed for TIE2 protein, which was significantly downregulated in the lungs of *Erg*^*iECko*^ mice but not the heart, liver, or kidney **(Fig. 5 E-F)**. These findings were further supported by immunostaining for TIE2 in lung sections collected from *Erg*^*iECko*^ mice, which revealed a substantial loss of TIE2 expression within CD31^+^ capillary ECs **(Fig. 5G)**. Therefore, ERG regulates *Tek* expression *in vivo* in an organotypic manner.

To determine if *Tek* is also regulated in an organotypic manner during a systemic state of inflammation, we performed intraperitoneal LPS injections in wildtype mice and used qPCR to assess *Tek* expression in the lung and heart at 6-72 h after LPS challenge. In both tissues, we observed a maximum *Tek* downregulation (>80%) at 6 h post-LPS injection. However, in the lung the downregulation of *Tek* expression persisted for up to 72 h, whereas in the heart *Tek* expression had recovered to a basal level by 24-48 h post-LPS injection **(Fig. 5H** and **I)**. Thus, *Tek* regulation after an LPS challenge *in vivo* in the lung vasculature mirrors that of ERG protein expression.

### EC-specific deletion of *Erg* leads to pulmonary vascular hyperpermeability and immune cell infiltration

ERG provides transcriptional support for vascular stability and barrier integrity through additional targets besides *TEK*^19,45,53,54^. We therefore turned our attention to understanding how the loss of ERG expression in pulmonary ECs might contribute to well-known features of the inflamed lung. First, we determined the impact of ERG downregulation on vascular permeability by performing an Evans blue dye leakage assay in *Erg*^*iECko*^ and littermate control mice. In agreement with a recent study^23^, lungs from *Erg*^*iECko*^ mice displayed increased pulmonary vascular permeability compared to those from control mice using fluorometric quantification of extravasated dye **(Fig. 6A**). However, we also noticed that ERG-dependent vascular hyperpermeability was more prominent in the lung when compared to the other organs tested. We therefore employed an alternative measure of vascular permeability by performing intravenous injection of FITC-Dextran (MW=150,000), which is too large to exit from most non-activated blood vessels. After one hour, mice were perfused to remove vascular FITC-Dextran, and lung, heart, liver, and kidney tissues were collected for analysis by immunostaining. We observed substantially more extravascular FITC-Dextran in the lungs of *Erg*^*iECko*^ mice compared to control mice, indicative of vascular hyperpermeability in the mutants **(Fig. 6B)**. In agreement with our Evans blue dye assay, deletion of endothelial *Erg* had a less dramatic effect on permeability in other organs **(Fig. 6C)**.

**FIGURE 6.**
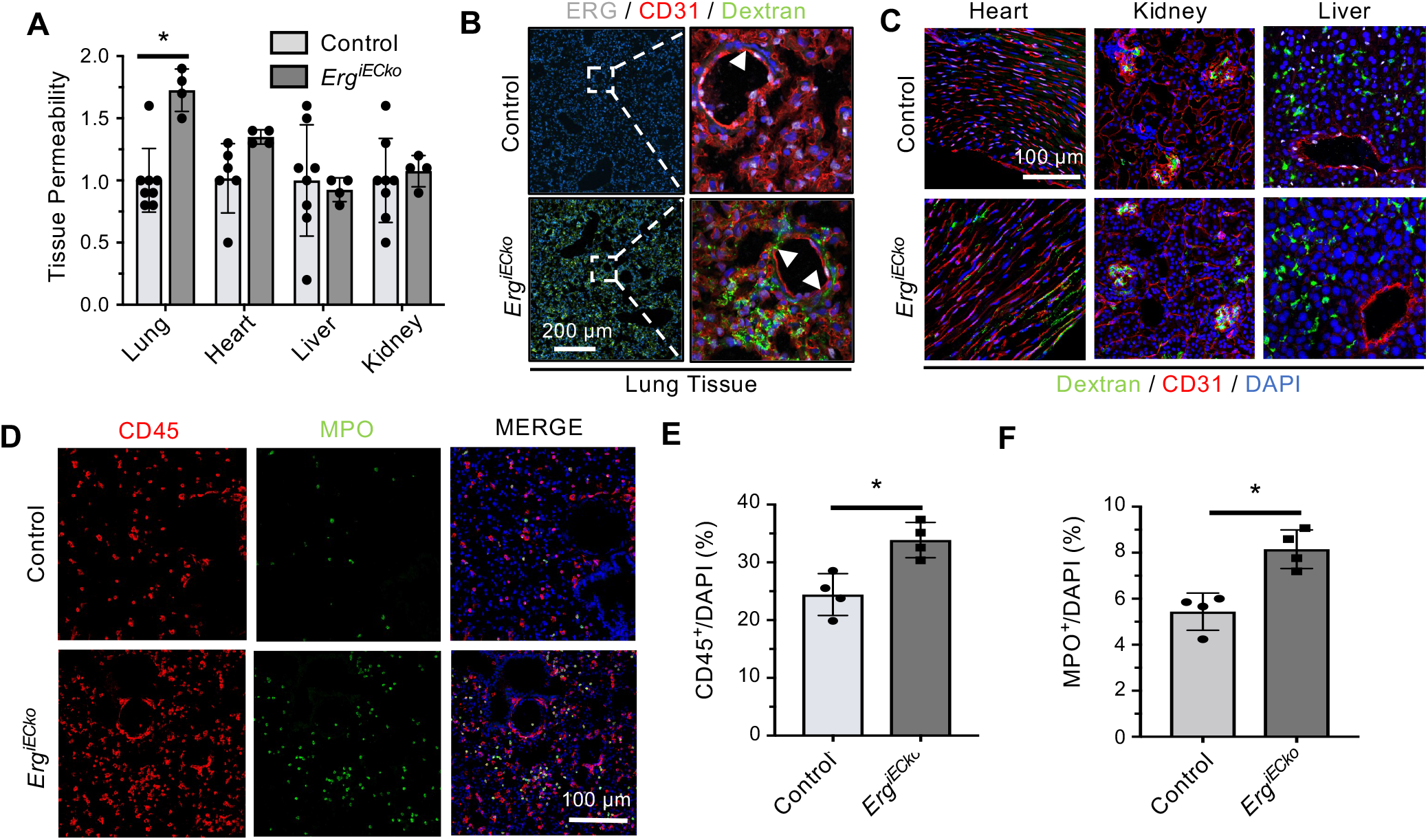
Vascular permeability and immune cell recruitment occurs in the lungs of Erg^iECko^ mice. **(A)** Evans blue dye leakage in the lung, heart, liver, and kidney from control and *Erg*^*iECko*^ mice was spectrophotometrically quantified and normalized to dry tissue weight. **(B**,**C)** Intravenously injected FITC-Dextran (green; MW=150,000 kDa) could be visualized outside of pulmonary blood vessels (CD31; red) (white arrows) in *Erg*^*iECko*^ mice but not in littermate controls (B). However, extravasation of FITC-Dextran in the heart, liver, and kidney following endothelial *Erg* deletion was less apparent (C) ERG (grey); DAPI (blue). **(D-F)** Greater numbers of CD45^+^ (red) immune cells and MPO^+^ (green) neutrophils were observed in the lungs of *Erg*^*iECko*^ mice relative to their control littermates (D), which was quantified by normalization against nuclei (DAPI; blue) (E-F). *p<0.05 (two-way t-test)

During inflammation, vascular hyperpermeability facilitates the recruitment and extravasation of immune cells into the lung and other tissues^55,56^. Moreover, ERG inhibition has been shown to promote immune cell recruitment *in vitro* and *in vivo*^20,23^. We therefore assessed the abundance of immune cells in the lungs of *Erg*^*iECko*^ and control mice using the pan-immune cell marker CD45^57^, revealing greater numbers of CD45^+^ cells in the lungs of *Erg*^*iECko*^ mice **(Fig. 6D-E)**. Importantly, a similar increase in pulmonary immune cells was also observed using an independently generated *Erg*^*iECko*^ line **(Supplemental Fig. 5B)**. Neutrophils are among the first immune cells recruited to the inflamed lung^58–60^. In agreement with others^23^, we observed a greater number of myeloperoxidase (MPO)-positive neutrophils in the lungs of *Erg*^*iECko*^ mice **(Fig. 6D** and **F)**. In contrast, *Erg* depletion had no apparent effect on the numbers of MPO^+^ immune cells in the heart, liver, or kidney **(Supplemental Fig. 5C-E)**.

### EC-specific deletion of *Erg* leads to organotypic tissue fibrosis and disrupts pulmonary vascular structure

Although vascular hyperpermeability and immune cell recruitment play beneficial roles in acute inflammatory responses, prolonged inflammation and vascular instability can lead to tissue fibrosis. We therefore used a Masson’s trichrome stain to assess collagen deposition in the lung, liver, heart, and kidney of control and *Erg*^*iECko*^ mice. Deletion of *Erg* resulted in a substantial increase in collagen deposition in the lung, accompanied by significant structural disruptions; the liver and heart also showed increased collagen deposition in *Erg*^*iECko*^ mice, albeit to a lesser extent, while no major difference was found for the kidney **(Fig. 7A)**. We therefore used transmission electron microscopy (TEM) to explore the consequences of *Erg* deletion on lung microvascular structures. In lungs from control mice, endothelial cells and pneumocytes form thin layers that separate the blood from alveolar air sacs. However, these structures were dramatically disrupted in *Erg*^*iECko*^ mice. We observed signs of EC hypertrophy and basement membrane delamination that led to a pronounced thickening of the membrane separating the blood and alveolar compartments **(Fig. 7B)**. Thus, our data indicate that despite its ubiquitous expression in ECs, the loss of ERG has organotypic functional and structural consequences that are most evident in the lung microvasculature.

**FIGURE 7.**
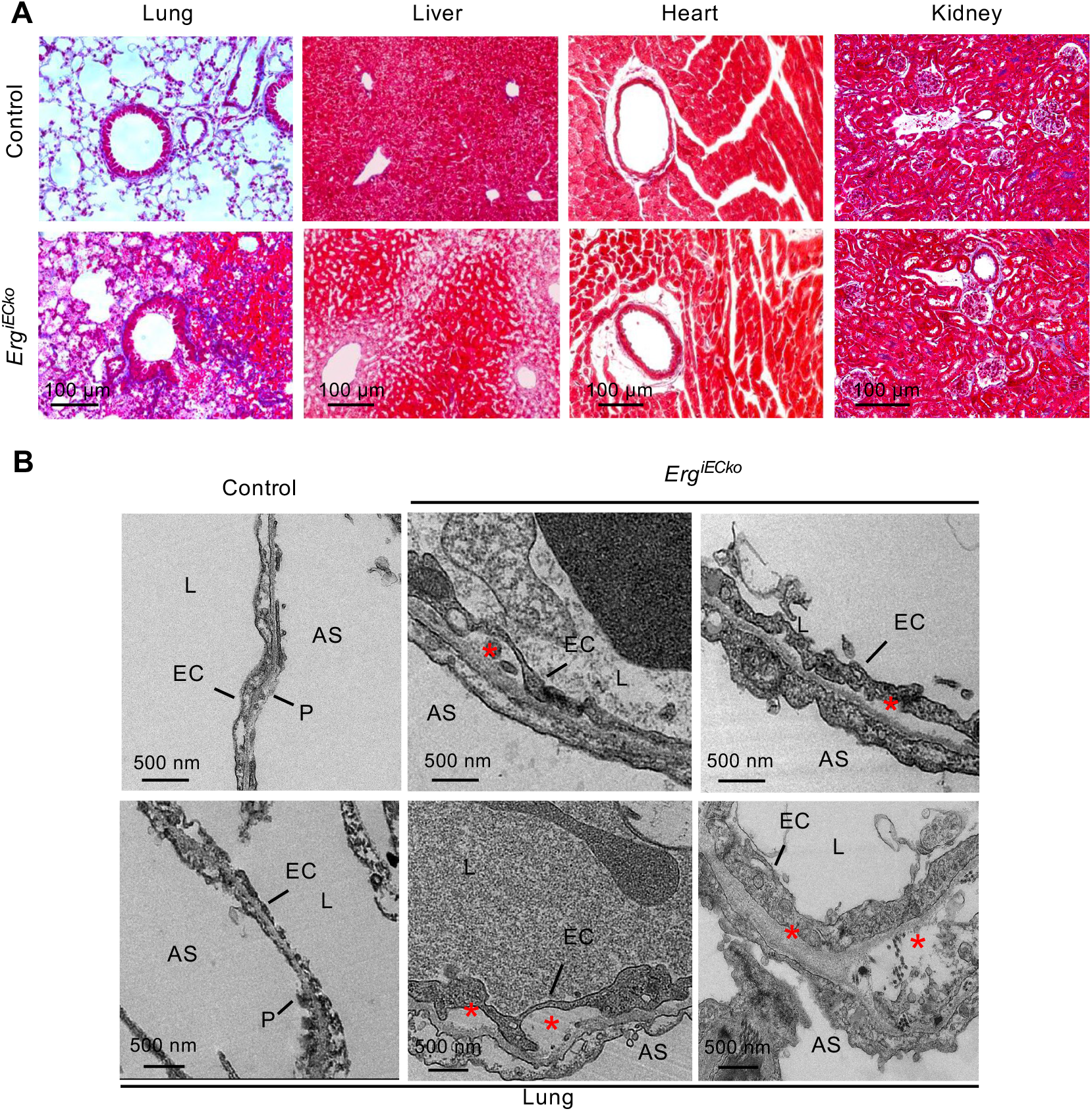
Deletion of endothelial Erg leads to organotypic fibrosis and disruption of vascular structures within the lung. **(A)** Masson’s trichrome staining was used to assess collagen (blue) deposition within the lung, liver, heart, and kidney of *Erg*^*iECko*^ and control mice. **(B**,**C)** Transmission electron microscopy was used to assess microvascular structures within the lung (B) and kidney (C) from *Erg*^*iECko*^ and control mice. In the lung, deletion of endothelial *Erg* led to EC hypertrophy and basement membrane delamination (red asterisks) resulting in a separation from pneumocytes (P) and the enlargement of the membrane separating alveolar air sacs (AS) and vascular lumens (L).

## DISCUSSION

Each year sepsis affects ∼20 million individuals worldwide with mortality rates among hospitalized patients ranging from 17-26%^61^. The progression of sepsis is complex, resulting from the collective disruption of multiple organ systems by a rampant pro-inflammatory response to bloodborne bacterial infection. However, impaired respiratory function is one of the most common manifestations of sepsis^62^. In 6-7% of patients, continued deterioration of pulmonary function results in ARDS for which mortality rates can be as high as 40%^63,64^. Within these patients, inflammation-induced destabilization of pulmonary blood vessels disrupts fluid homeostasis and impairs efficient gas exchange between the blood and alveolar air sacs thereby undermining organ function^65^. In addition to sepsis, ARDS can result from seemingly diverse conditions including viral/fungal infection^66,67^, burn injury^68^, brain injury^69^, drug overdose^70^, or pancreatitis^71^, suggesting that the lung may be generally prone to vascular dysfunction under inflammatory challenge. Yet the mechanistic reasons for this predisposition are poorly understood. As a result, we currently lack therapeutic options that effectively target the etiology of vascular dysfunction within the lungs of sepsis/ARDS patients^65^.

Here we have demonstrated how organotypic regulation of the master endothelial transcription factor ERG preferentially destabilizes pulmonary blood vessels during a systemic inflammatory challenge. Acute LPS exposure led to the downregulation of *Erg* mRNA transcripts in the lung, heart, liver, and kidney, yet the rapid downregulation of ERG protein was only observed in the lung. We speculate that these are independent processes collectively resulting in a more rapid and substantial loss of ERG protein in the lung when compared with other organs. Although ERG proteolysis has primarily been studied in the context of cancerous prostate epithelial cells, in which oncogenic ERG fusion proteins evade recognition by the SPOP and TRIM25 ubiquitin ligase^72,73^, two recent studies have documented ERG degradation in cultured ECs^24,25^. Here we have generated a list of ubiquitin ligases expressed in ECs from multiple organs to ask if lung ECs harbor unique proteolytic that may target ERG for ubiquitination/degradation. Notably, this list confirms that both TRIM25 and SPOP are expressed by pulmonary ECs suggesting that they may similarly regulate ERG degradation in vivo. However, given that their expression is not confined to the lung, other mechanisms must be involved in the control of organotypic ERG degradation. We identified an additional ∼30 ubiquitin ligase that were >2-fold higher expressed in lung ECs compared to other organs. Interestingly, many of these ubiquitin ligases have not yet been studied in the context of EC function. Therefore, the underlying mechanism for lung-specific ERG degradation remains unclear and will require further study.

The transcriptional responses of activated ECs vary across vascular beds^40,74^. Therefore, lung-specific degradation of ERG, which has well documented anti-inflammatory and pro-homeostatic functions, likely contributes to organotypic transcriptional responses of pulmonary ECs^19^. Indeed, loss of endothelial *Erg* expression has a substantial effect on pulmonary vascular function, as recently demonstrated in a murine model of the aging lung^23^. In agreement with that study, we found that inducible genetic deletion of endothelial *Erg* is sufficient to drive pulmonary vascular permeability, immune cell recruitment, and fibrosis. However, our study brings new information to light by demonstrating that these effects were more dramatic in the lung despite the deletion of *Erg* in all ECs, suggesting that ERG may have organotypic functions within distinct vascular beds.

In support of this hypothesis, we found that deletion of endothelial *Erg* predominantly downregulated *Tek*/TIE2 expression within the lung. Reduced expression of *Tek*/TIE2 plays a causal role in the destabilization of pulmonary blood vessels during sepsis/ARDS^47,49^. Our data indicate that following acute LPS challenge, *Tek* expression in the lung and heart is initially downregulated to a similar extent, yet basal *Tek* expression was restored more slowly in the lung. Interestingly, the recovery of *Tek* transcripts coincided with the recovery of ERG protein expression following acute LPS challenge. Therefore, we speculate that within the inflamed lung the loss of ERG expression impairs the recovery of *Tek* expression following its downregulation during the acute phase of inflammation. Importantly, ERG regulates many additional genes that influence the vascular inflammatory response^20,21,37,53,54,75^. Therefore, it will be interesting to determine if these genes are likewise regulated by ERG in an organotypic and stimulus-specific manner.

Constitutive deletion of *Erg*, which is highly expressed by nearly all ECs, causes embryonic lethality between day E10.5 and E11.5^32^. Postnatal downregulation of endothelial ERG is associated with multiple organ phenotypes, including tissue fibrosis^23,76^, bleeding and thrombosis^75^, and ocular vascular regression^77^ via the regulation of critically important pathways such as Wnt/β-Catenin^32^, TGFβ^76^, KLF-2^75^ and angiopoietin signaling^18^. Accordingly, loss of ERG expression in human diseases is associated with atherosclerosis^54^, liver fibrosis^76^, and pulmonary hypertension^22^. For these reasons, ERG has been identified as a critical mediator of vascular function^19^. ERG is frequently co-expressed in ECs along with its homologue FLI1, with which it shares ∼75% sequence similarity. ERG and FLI1 exhibit overlapping and often compensatory functions^22,78,79^, and we recently demonstrated loss of vascular identity and systemic vascular collapse following the inducible endothelial deletion of both *Erg* and *Fli1* in adult mice^39^. Therefore, our observation that both ERG and FLI1 are rapidly downregulated in the lungs of LPS-challenged mice suggests a common mechanism to their degradation and raises questions about their combinatorial impact on the response to vascular inflammation in pulmonary ECs.

Altogether, this study adds to our growing appreciation of EC heterogeneity by providing new insights into the organotypic vascular response to inflammation. Our work indicates that the widely expressed endothelial transcription factor ERG, a master regulator of endothelial function, is targeted by cytokines for preferential proteolytic degradation in the lung. Based on our data, we propose that during inflammation acute ERG downregulation helps coordinate a transcriptional response within pulmonary ECs to support vascular hyperpermeability and immune cell recruitment. However, sustained loss of endothelial ERG impairs the ability of the lung to return to homeostasis, thereby predisposing the lung to vascular dysfunction. Therefore, identification of the mechanism by which inflammatory cytokines trigger pulmonary ERG degradation could yield new therapeutic targets for combating lung-damaging diseases such as sepsis and ARDS.

## AUTHOR CONTRIBUTIONS

CS, SMA, GB, AR, and CG designed the study and interpreted the results. CS, SMA, KK, ND, LOA, MW, CF, AS, AB, EW, and KW, performed the experiments and data analysis and contributed to the interpretation of results. ST, MC, and SK performed the intratracheal influenza infections. RS performed and interpreted the TEM. DH contributed to the study design and edited the manuscript. CS wrote the manuscript with the help of AR and CG. All other authors edited the manuscript.

## ACKNOWLEDGEMENTS

We would like to thank Professor Justin C. Mason (Imperial College London) for his constant support and advice. We would also like to thank Jun Xie for mouse colony management and the C.T.G. laboratory members for helpful discussions.

